# Investigation of polymorphisms in the *P. falciparum* artemisinin resistance marker *kelch13* in asymptomatic infections in a rural area of Cameroon

**DOI:** 10.1101/148999

**Authors:** Innocent Safeukui, Jerome Fru-Cho, Alassane Mbengue, Niraja Suresh, Dieudonne L. Njimoh, Violet V Bumah, Theresa Nkuo-Akenji, Vincent PK Titanji, Kasturi Haldar

**Affiliations:** Boler-Parseghian Center for Rare and Neglected Diseases, University of Notre Dame, 103 Galvin Life Sciences, Notre Dame, IN 46556, USA.; Department of Biological Sciences, University of Notre Dame, 103 Galvin Life Sciences, Notre Dame, IN 46556, USA.; Department of Microbiology and Parasitology, University of Buea, Buea, South West Region, Cameroon; Department of Biochemistry and Molecular Biology, Faculty of Science, University of Buea, P.O. Box 63 Buea, Southwest Region, Cameroon; Department of Biology, North Life Science 317, San Diego State University, CA 92182, USA

**Keywords:** Asymptomatic malaria, Plasmodium falciparum, Kelch 13 polymorphisms, artemisinin resistance

## Abstract

**Background:** The genetic variability of the artemisinin resistance (AR) molecular marker *kelch13 (k13)* has been extensively investigated in *Plasmodium falciparum* malaria parasites from symptomatic infections in South East (SE) Asia where AR is highly prevalent, as well as in Africa where evidence of AR has emerged only recently. However, molecular surveillance and risk of transmission of AR also require monitoring asymptomatic infection. Here, molecular analyses were used to investigate polymorphisms in *k13* and their potential for transmission in asymptomatic adults in Bolifamba, Cameroon in Central Africa.

**Methods:** Using polymerase chain reaction (PCR), we amplified and sequenced the full length of *k13* from *P. falciparum* infections detected in the blood of 33 asymptomatic adults (age: 18-55 years-old) collected in a cross-sectional study from July 2008 to October 2009. Risk of increased transmission was assessed by quantifying gametocytes by qPCR. Quantitative ELISA was used to detect plasma levels of PfHRP2 to establish total parasite burdens associated with asymptomatic infection.

**Results:** Out of 33 isolates tested, 14 (42.4%) presented at least one single nucleotide polymorphism (SNP) in *k13.* Five non-synonymous SNPs were detected (K189T/N, N217H, R393K and E433K). None were located in the ß-propeller domain, where AR mutations have been detected in both SE Asian and, more recently, African parasites. K189T/N and N217H have been previously reported in African strains, but R393K and E433K are new polymorphisms. Gametocytes were detected in 24.2% of infections, without significant association with detected *k13* polymorphisms. Notably, polymorphisms outside of the ß-propeller domain detected in *k13* were associated with a significant increase of PfHRP2 plasma levels but not circulating parasite levels detected by qPCR.

**Conclusions:** This study provides the baseline prevalence of *k13* polymorphisms in asymptomatic infection for molecular surveillance in tracking AR. Unexpectedly, it also suggests association of *k13* polymorphisms outside of the ß-propeller domain with total *P. falciparum* burden in asymptomatic infection, that needs to be validated in future studies.

## Background

Artemisinin combination therapy (ACT), the frontline treatment for *Plasmodium falciparum* infection, has made a major contribution in reducing worldwide malaria burden and deaths over the past decate [1]. However, the appearance of the artemisinin resistant (AR) *P. falciparum* in western Cambodia [2] and its emergence and spread throughout South East (SE) Asia threatens efforts to control and eliminate malaria [3,4].

The *P. falciparum k13* gene has been identified as a molecular marker of AR [5], and two major molecular mechanisms have emerged [6,7]. Single nucleotide polymorphisms (SNPs) situated in the β-propeller domain of *k13* associated with delayed parasite clearance in patients and, characteristic of clinical AR in SE Asia have been validated in the ring-stage survival assay (RSA) [3,5], an *in vitro* correlate of clinical resistance [8]. The most prevalent clinical mutation C580Y causes resistance across parasite strains and is associated with 85% of the artemisinin treatment failure in SE Asia [5,9]. C580Y has been detected in African isolates at a very low frequency (0.06 - 2.7% of samples) [10 – 12] but not shown to be associated with clinical resistance in Africa [13]. A578S is the most common and frequent *k13* propeller polymorphism in Africa, but this mutation is not associated with clinical or *in vitro* AR [12 – 14]. Recently, a single case of clinical resistance associated with M579I in a febrile traveller returning from Equatorial Guinea (in Central Africa) was confirmed by *in vitro* RSA [15]. But AR is not widespread and ACT treatments remain widely effective in Sub-Saharan Africa [1]. Since 90% of *P. falciparum* malaria cases occur in Africa [1], a major current global health challenge is to prevent widespread dissemination of AR parasites throughout Africa. To accomplish this, it will be necessary to strengthen *P. falciparum* genotypic resistance surveillance in both symptomatic and asymptomatic infections since both can contribute to parasite transmission.

*k13* polymorphisms associated with AR are well documented in symptomatic *P. falciparum* infections [2, 3, 5, 9, 12]. However, only a few studies (almost all from SE Asia) document the presence of AR *k13* mutations in asymptomatic infections [16, 17]. This suggests that asymptomatic infections may represent an important reservoir for parasite dissemination [3]. Additionally, new *k13* polymorphisms in African strains may confer AR since the extent of resistance conferred is also determined by parasite strain background [18, 19]. Finally, historical information on *k13* polymorphisms provides important information on their emergence in the context of the time line of adoption of ACT as the national antimalarial drug policy. In this study, we examined the genetic variability of *k13* in asymptomatic *P. falciparum* infections in Cameroon (in 2008), four years after artesunate-lumefantrine (AL) was adopted as the first-line treatment of uncomplicated malaria in 2004. We also examined the potential effects on parasite transmission and total parasite load by quantifying respectively peripheral gametocytes and plasma levels of PfHRP2.

## Methods

### Ethics statement

All subjects gave signed informed consent before enrollment into the study. The research was approved by the University of Buea Institutional Review Board (IRB) and the IRB of the University of Notre Dame.

### Patient study design and sample collection

Samples analyzed in this study are from a cross-sectional study conducted from July 2008 to October 2009 in Bolifamba in the South West Region of Cameroon. The study design for sample collection from non febrile, asymptomatic patients and their rates of infection by *P. falciparum* as well as *P. vivax* has been previously described [20]. The experimental design summarized in Figure 1 provides a brief summary of the sample collections, where a total of 269 afebrile subjects, aged 18 to 55 year, residents of Bolifamba were enrolled. Indicated clinical parameters were recorded, blood samples were collected in EDTA tubes and the plasma was separated by centrifugation (1,000 × g, 10 min), aliquoted and frozen at −80°C. Eighty-seven out of 269 participants were positive for *Plasmodium* spp. as determined by polymerase chain reaction (PCR) targetting the 18S ribosomal RNA gene [20]: Sixty five were positive only for *P. falciparum,* and 22 were positive for other *Plasmodium* species or mixed infection *(P. falciparum* + other species).

**Figure 1:**
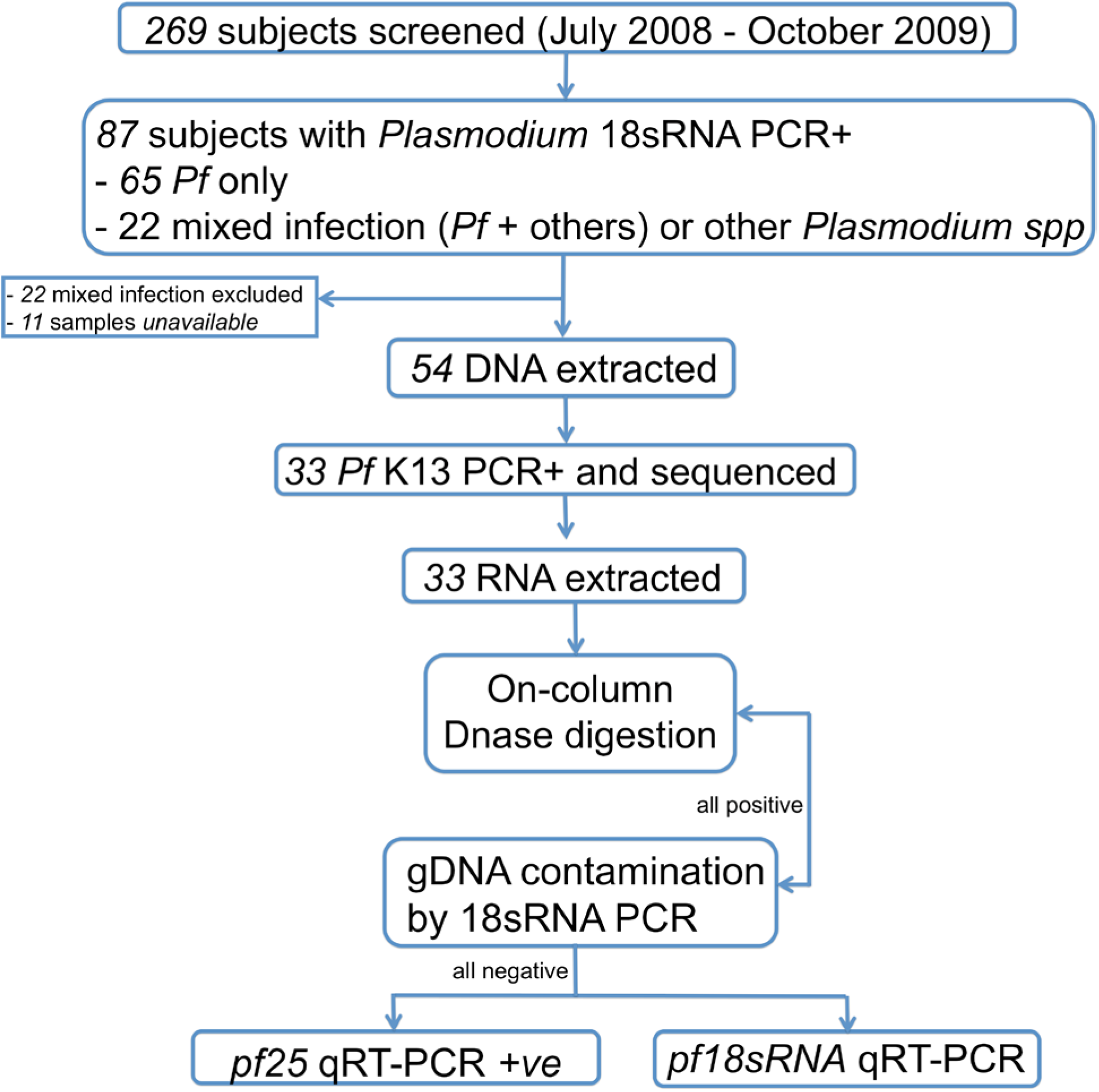
Experimental Design.

### DNA and RNA extraction

Total nucleic acid was extracted from 200 μL of blood pellet using Quick-gDNA Blood Miniprep kit (reference no. D3072, Zymo Research) (for total DNA) and RNeasy Lipid Tissue Mini kit (reference no. 74804, Qiagen) (for total RNA) according to the manufacturer’s instructions. Nucleic acid extracts were quantified using a NanoDrop ND-1000 (NanoDrop Technologies, Wilmington, DE, USA), and the samples immediately stored at −20°C until used.

### Genotyping

In order to identify the mutations in *k13,* the full length gene was amplified by PCR as previously described [7]. Briefly, the *k13* PCR was performed in a final volume of 50μL containing 50 to 100 ng of gDNA, 5 μL of Takara PCR buffer (Takara Bio Inc.), 4μL dNTPs mixture (10mM of each dNTP), 1μL MgCl2 (25mM), 4μL of the primers mixture (10 mM each forward and reverse), 0.5 μL of Takara Ex Taq Hot Start version (Takara Bio Inc.) and sterile autoclaved ultrapure water. The PCR reaction conditions and primers are listed in Supplemental Table 1. The PCR products were analyzed using 1% agarose gel electrophoresis. The *k13* DNA fragment was purified on agarose gel using NucleoSpin® Gel and PCR Clean-up kit (catalog reference no 740609.50, Macherey-Nagel Inc.) according to the manufacturer’s instructions, and sequenced. The primers used for sequencing are presented in Supplemental Table 1. The 3D7 *k13* sequence was used as reference.

### Quantitative real-time PCR

The asexual and sexual stages of *P. falciparum* were quantified by quantitative real-time PCR (qPCR) targetting 18S ribosomal RNA gene transcripts (M19173.1, for asexual stages) or transcripts specific for mature gametocytes (PF10_0303; Pbs25). The qPCR was performed using the 7900 HT Fast real-time PCR system (Applied Biosystems) with a 20 μL reaction volume and the Power SYBR Green RNA-to-CT one-step kit (catalog reference no. 4389986; Applied Biosystems). The qPCR reaction conditions and primers are listed in Supplemental Table 1. As a positive control to ensure parasite RNA was present in our samples, we used primer pairs for *P. falciparum* ubiquitin conjugating enzyme transcript (PF08_0085) as a constitutively expressed parasite marker as previously described [21]. For asexual stages, RNA extracted from parasite cultures (strain NF54) with different parasitemia (as determined by Giemsa-stained blood smears) was used to develop standard curves. Each RNA sample was run in duplicate.

### Quantitative PfHRP2 measurements

The quantification of *P. falciparum* histidine-rich protein 2 (PfHRP2) in the plasma of malaria patients was carried out by sandwich ELISA as previously described [22]. The mouse monoclonal antibodies anti-P. *falciparum* HRP2 IgM (MPFM-55A; Immunology Consultants Laboratory Inc., USA) and horseradish peroxidase (HRP)-conjugated anti-*P. falciparum* IgG (MPFG-55P; Immunology Consultants Laboratory Inc., USA) were used for plate coating and detection, respectively. The detection limit of the assay was 31.25 pg/mL. Positive cases were defined as those in which duplicate derived concentrations were greater than the detection limit. Samples were analyzed blinded with respect to to *k13* polymorphisms.

### Statistical analysis

The number of gametocyte carriers were expressed as a percentage of isolates harboring mature gametocytes among groups of isolates presenting at least one SNP *(k13* polymorphism+) or without SNP *(k13* polymorphism-). For plasma concentrations of PfHRP2 and peripheral parasitemia, we used median and interquartile range (IQR) or geometric mean and 95% confidence interval (95% CI). Chi-square test was used to compare the proportions of isolates positive for gametocytes and the non-parametric Mann-Whitney U test was used to compare PfHRP2 concentrations and parasitemia. The statistical analysis was performed with GraphPad Prism (version 6.02). p-values were two sided, with p <0.05 being considered significant.

## Results

In this study, DNA was extracted from 54 isolates with *P. falciparum* mono-infection collected from asymptomatic adults. The full length of *k13* gene was successfully amplified by PCR and sequenced from 33 isolates only. Table 1 summarizes the baseline (day 0) of the clinical and laboratory characteristics of the 33 included subjects. Twenty two individuals were females and 11 were males. The median [IQR] of the participant parameters were as follows: age 24 [20 – 30] years; axillary temperature 36.8 [36.3 - 37.1]°C; parasitemia 0.15 [0.11 - 0.29] parasite/μL; hemoglobin concentration 12.3 [11.8 - 13.9] g/dL and packed cell volume 39 [35–42] %.

**Table 1:** The baseline (day 0) of the clinical and laboratory characteristics of participants from whom DNA could be extracted and the full length of *k13* amplified by PCR and sequenced.

| Parameters | Day0 |
| --- | --- |
| Age (years) | 24 (20 - 30) |
| Gender (female/male) | 22/11 |
| Body temperature (°C) | 36.8 (36.3 - 37.1) |
| Hemoglobin (g/dL) | 12.3 (11.8 - 13.9) |
| Pack cell volume (%) | 39 (35 - 42) |
| *P. falciparum* 18sRNA or pfs25 qRT-PCR |  |
| Parasitemia (parasite/μL) | 0.15 (0.11 - 0.29) |
| Gametocytes, n(%) | 8 (24.2) |
| Plasma PfHRP2 (pg/mL) | 482.4 (60.9 - 2289.7) |
| *k13* polymorphisms, n(%) |  |
| K189T | 7 (21.2) |
| K189N | 1 (3.0) |
| N217H | 2 (6.1) |
| R393K* | 1 (3.0) |
| E433K* | 5 (15.2) |

Values are median (interquartile range); n, number of patients; *polymorphism not previously reported

The genotyping of *k13* revealed five different SNPs (Table 1). As observed in many African countries [12 – 14, 23, 24], the *k13* was highly polymorphic: 14 out of 33 (42.4%) of the tested isolates present at least one SNP in the BTB/POZ domain of the protein. Three SNPs (K189T, K189N and N217K) have been previously described [3, 23, 24] and two SNPs (R393K and E433K) have not been reported (Table 1). For most of the isolates, the polymorphisms occurred singly, except for two isolates with double-SNPs (K189T/E433K and N217H/R393K). The K189T substitution (highly prevalent in African samples) was the most frequent (21.2%) followed by the E433K mutation (15.2%) (Table 1).

Since asymptomatic infection is critical for the control of malaria as a transmission reservoir of the parasites [16, 17], we quantified mature gametocytes by qPCR targetting their specific transcripts (PF10_0303; Pbs25). The gene selection was based on previous studies showing their expression as a specific marker for mature gametocytes [21]. We found that 24.2% of these isolates contain mature gametocytes (Table 1). This prevalence was 35.7% in isolates presenting at least one SNP (*k13* polymorphism+) compared to 15.8% in isolates without SNP *(k13* polymorphism-), although the difference was not statistically significant (p > 0.05). This is probably due to the low sample number. There was no correlation with peripheral parasitemia quantified by qPCR (Figure 2).

**Figure 2:**
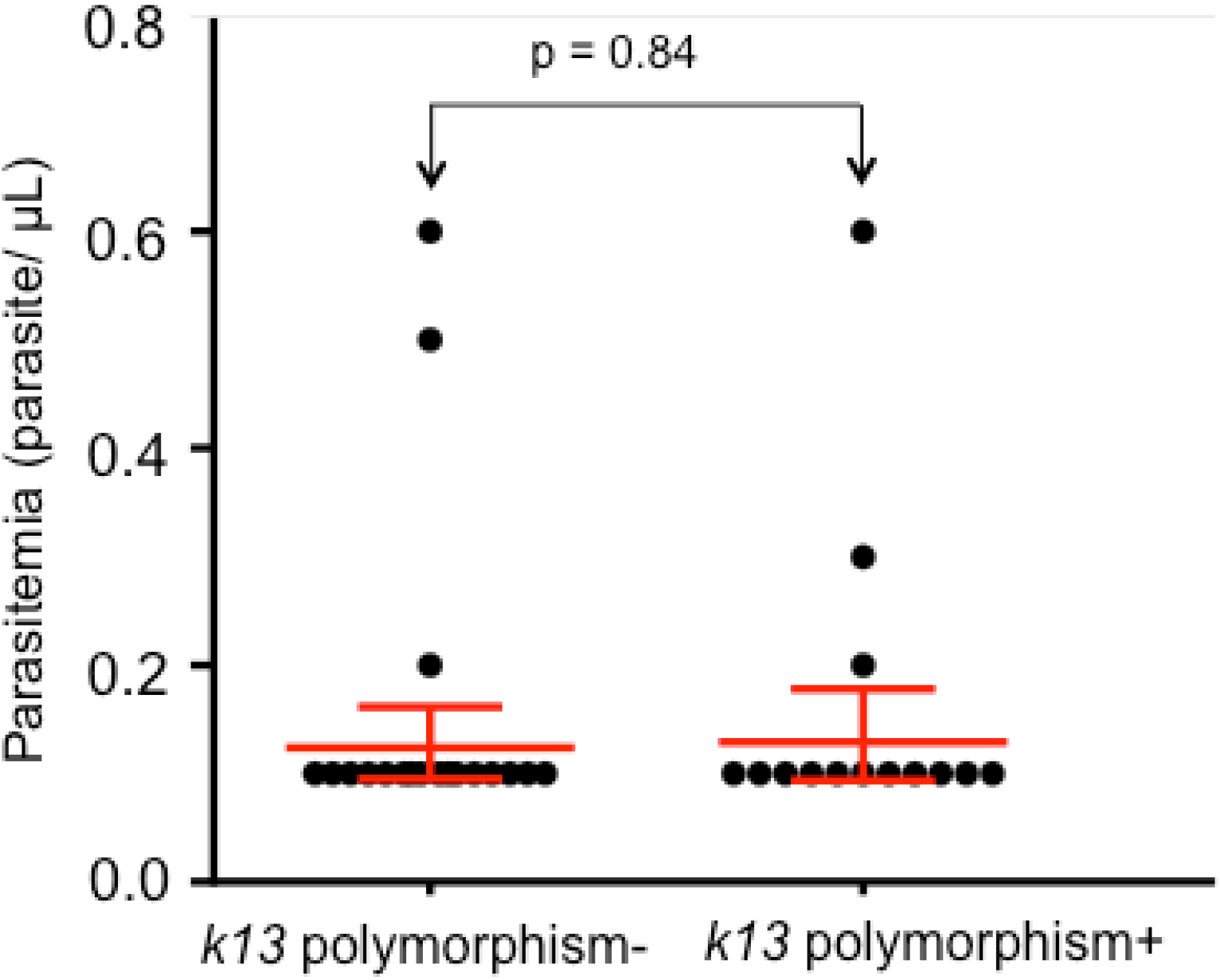
*k13* polymorphism and peripheral parasitemia quantified by qPCR. The horizontal bar represents the geometric mean value and the vertical bar 95% CI. Mann-Whitney U test was used to compare the mean of parasitemia between individuals with isolates presenting at least one *k13* SNP *(k13* polymorphism+, n = 14) and without SNP *(k13* polymorphism-, n = 19).

PfHRP2, the lead marker used to estimate total parasite mass [25], was detected in over 79% of patient samples (13 out of 14 in *k13* polymorphism+ and 13 out of 19 in *k13* polymorphism-). The median [IQR] values of PfHRP2 concentrations were significantly higher in samples with *k13* polymorphism (1500.8 [419.1 - 8803.6]) compared to those of without *k13* polymorphism (323.0 [6.5 - 797.3]) isolates (p = 0.024) (Figure 3).

**Figure 3:**
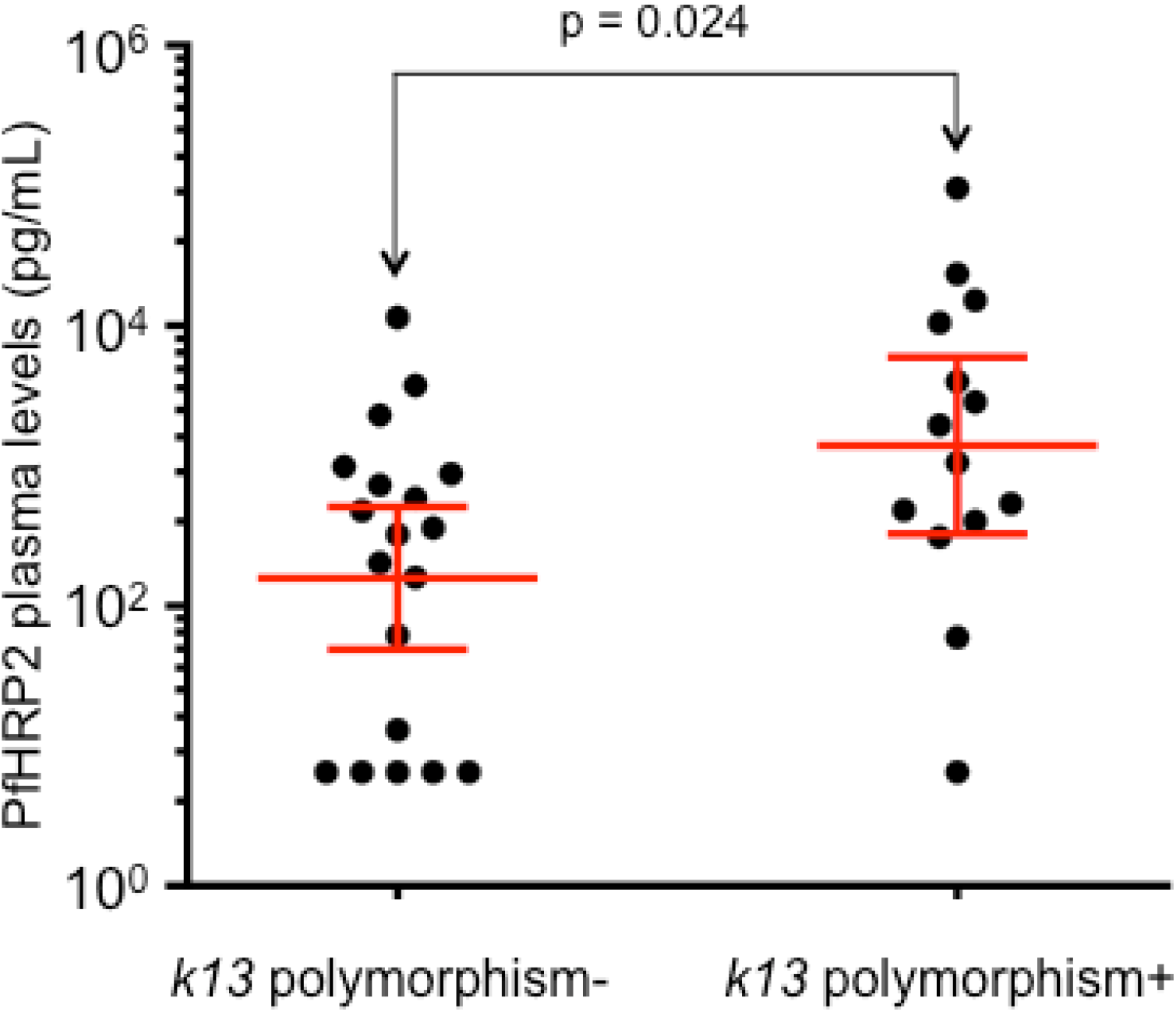
Plasma levels of PfHRP2 from individuals with isolates presenting at least one *k13* SNP *(k13* polymorphism+, n = 14) or without SNP *(k13* polymorphism-, n = 19). The horizontal bar represents the geometric mean value and the vertical bar 95% CI. Isolates with undetectable PfHRP2 by ELISA (one isolate in *k13* polymorphism+ and 6 isolates in *k13* polymorphism-) were given the value of the detection limit of the assay (31.25 pg/mL) divided by two. Mann-Whitney U test was used to compare plasma concentrations of PfHRP2 between the two groups.

## Discussion

This study investigated the genetic variability of *k13,* the molecular marker of AR in asymptomatic *P. falciparum* infections in the Bolifamba Southwestern Cameroon in 2008. This was four years after AL was adopted as the first-line treatment of uncomplicated malaria in 2004. As observed in many African countries [12, 14, 23, 24], *k13* was highly polymorphic as 42.4% of the isolates analyzed present at least one SNP. The K189T substitution, frequent in African isolates [3, 23, 24], was the most common (21.2%), although at a lower frequency compared to that observed in other African countries such as Uganda (34.4%) [23] and Senegal (42.2%) [24].

We did not find any polymorphisms in the *k13* propeller domain where mutations have been associated with AR in SE Asia [3, 5], consistent with the efficacy of ACT in Cameroon [1, 11]. Recent studies with *P. falciparum-infected* isolates collected from febrile returning travellers and migrants, or residents from Cameroon revealed the presence of SNPs in the propeller domain of *k13* [11 – 13], none of which were associated with artemisinin treatment failure or *in vitro* AR [11, 13, 26].

The absence of ACT treatment failure observed in African sites could be due to the presence of a strong “premunition” (partial immunity acquired during repeated infections throughout life) which potentiates the action of antimalarial drugs, and also to the very low fraction of the mutant parasite populations. Nonetheless the massive use of ACT in Africa may give transmission advantage to AR *P. falciparum* infections to drive the spread of resistance. Although we found that 35.7% of the isolates containing at least one SNP harboured mature gametocytes our sample size was insufficient to determine significance (and none of the mutations are associated with AR).

We found that the presence of SNP in *k13* was associated with higher plasma concentrations of PfHRP2 (but not peripheral parasitemia). We could not analyze the effect of a specific SNP due to the low number of tested samples. The molecular basis of this association is unknown. This observation stresses the need for additional studies to explore the association of *k13* polymorphisms with *P. falciparum* burden independent of AR, in this region of Cameroon.

## Conclusions

This study provides the baseline prevalence of *k13* polymorphisms in asymptomatic infection for molecular surveillance in tracking AR. It also suggests association of *k13* polymorphisms outside of the β-propeller domain with total *P. falciparum* burden in asymptomatic infection, that needs to be validated in future studies.

## Abbreviations

AR: artemisinin resistance
SE: South East
PCR: polymerase chain reaction
ELISA: enzyme-linked immunosorbent assay
qPCR: quatitative polymerase chain reaction
PfHRP2: *Plasmodium falciparum* histidine-rich protein 2
SNP: single nucleotide polymorphism
ACT: Artemisinin combination therapy
k13: kelch 13
RSA: ring-stage survival assays
AL: artesunate-lumefantrine
IRB: Institutional Review Board
EDTA: Ethylene diamine tetraacetate
DNA: Deoxyribonucleic acid
RNA: ribonucleic acid
dNTP: Deoxynucleotide triphosphate
IQR: interquartile range 95%
CI: 95% confidence interval

## Declarations

### Authors’ contributions

IS designed the study, performed experiment, analyzed and interpreted the data, wrote the manuscript, and has given final approval of the version to be published. JF-C designed the study, performed experiment, analyzed and interpreted the data, revised the manuscript, and has given final approval of the version to be published. AM designed the study, analyzed and interpreted the data, revised the manuscript and has given final approval of the version to be published. NS designed the study, analyzed and interpreted the data, revised the manuscript and has given final approval of the version to be published. DLN designed the study, revised the manuscript and has given final approval of the version to be published. BVV designed the study, revised the manuscript and has given final approval of the version to be published. TN-A supervised the study and also revised the manuscript, and has given final approval of the version to be published. VPKT designed and supervised the study, analyzed the data and also revised the manuscript, and has given final approval of the version to be published. KH designed the study, supervised the study, analyzed and interpreted the data, revised the manuscript, and has given final approval of the version to be published.

## Acknowledgements

This work was supported in part by grants to KH (NIH R01 HL069630, R01 HL130330 and the University of Notre Dame), TN-A (UNDP/ World Bank/ WHO Special programme for Research and Training in Tropical Diseases Grant No.990965) and VPKT (The International Program in the Chemical Sciences (IPICS) – CAM 01 project, and Microsoft Corporation).

---

**Supplemental Table 1.**
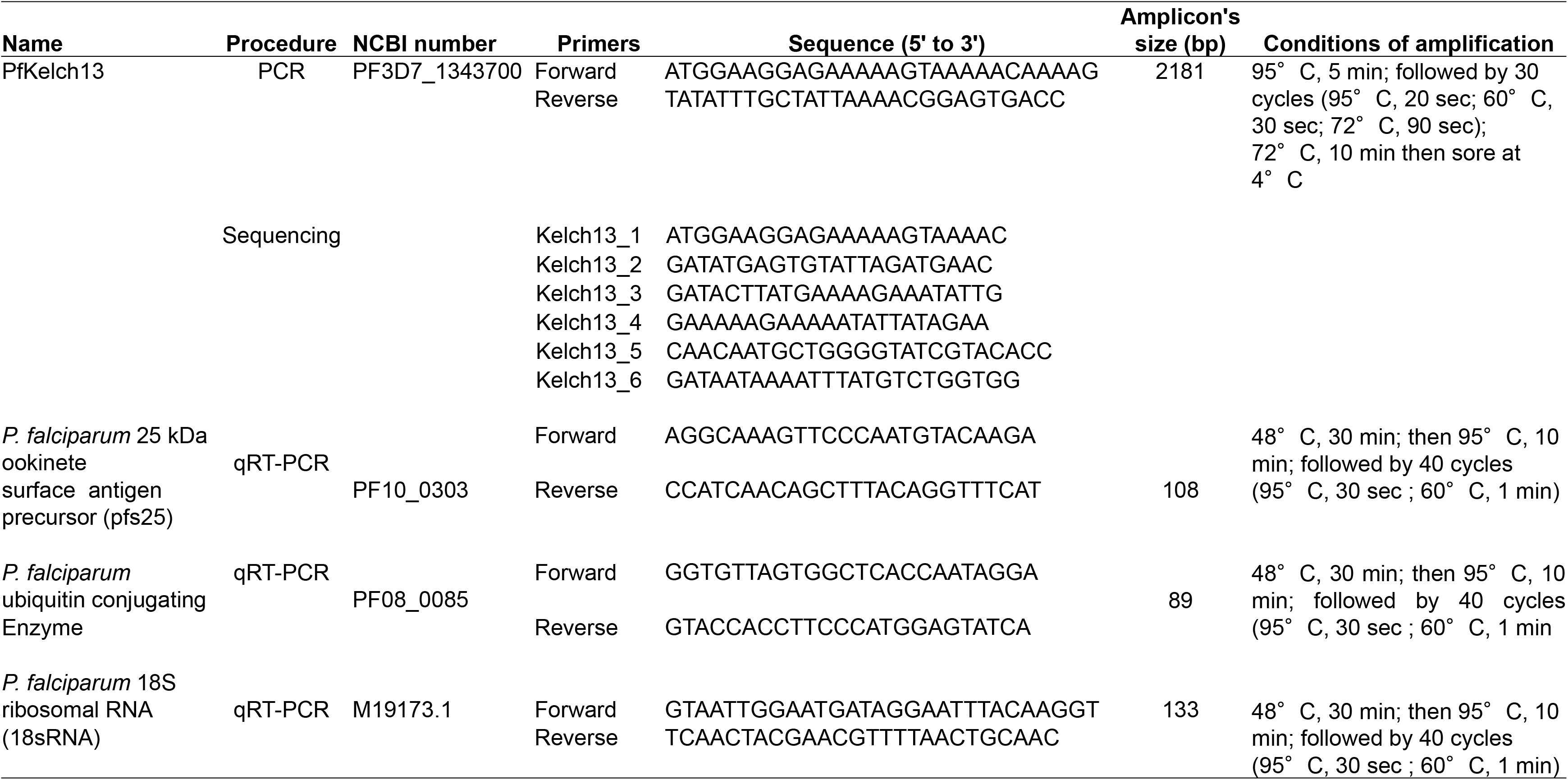
PCR and qRT-PCR primers, and conditions.

